# Sensitivity and validation of porous membrane electrical cell substrate impedance spectroscopy (PM-ECIS) for measuring endothelial barrier properties

**DOI:** 10.1101/2023.07.11.548631

**Authors:** Alisa Ugodnikov, Oleg Chebotarev, Henrik Persson, Craig A. Simmons

**Affiliations:** Translational Biology & Engineering Program, Ted Rogers Centre for Heart Research, Toronto, ON Canada M5G 1M1; Institute of Biomedical Engineering, University of Toronto, Toronto, ON Canada M5S 3G9; Department of Mechanical & Industrial Engineering, University of Toronto, Toronto, ON Canada M5S 3G8

**Keywords:** impedance spectroscopy, biosensors, ECIS, barrier function, TEER, cell culture insert

## Abstract

Conventional trans-endothelial electrical resistance (TEER) setups are invasive and cannot directly measure monolayer integrity in co-culture. These limitations are addressed by porous membrane electrical cell-substrate impedance sensing (PM-ECIS), which measures barrier integrity in cell monolayers grown directly on permeable membranes patterned with electrodes. Here we advance the design and utility of PM-ECIS by investigating its sensitivity to working electrode size and correlation to TEER. Gold electrodes were fabricated on porous membrane inserts using hot embossing and UV lithography, with working electrode diameters of 250, 500, and 750 µm within the same insert. Frequency scans of confluent primary endothelial monolayers showed normalized resistance peaking at 4 kHz, consistent with traditional solid substrate ECIS. Sensitivity to resistance changes (4 kHz) during endothelial barrier formation was inversely proportional to electrode size, with the smallest electrodes being the most sensitive (p<0.001). Similarly, smaller electrodes were most sensitive to changes in impedance (40 kHz) corresponding to cell spreading and proliferation (p<0.001). Barrier disruption with EGTA was detectable by 250 and 750 µm (p<0.01), and 500 µm electrodes (p=0.058). Resistances measured by PM-ECIS vs. TEER for sodium chloride solutions were positively and significantly correlated for all electrode sizes (r>0.9; p<0.0001), but only with 750 µm electrodes for endothelial monolayers (r=0.71; p=0.058). These data inform design and selection of PM-ECIS electrodes for specific applications, and support PM-ECIS as a promising alternative to conventional TEER due to its capacity for direct, non-invasive, and real-time assessment of cells cultured on porous membranes.

## 1. Introduction

Biological barriers, which separate body compartments and the external environment, play key roles in homeostasis and protection of the body, through regulating nutrient transport, maintaining concentration gradients, and excluding pathogens [1]. The structural and functional role of biological barriers is accomplished through tissue-specific combinations of endothelial or epithelial cells, junctional proteins, transporters, supporting cells and matrices [2]. Disruption of these components is associated with various disease states, including cancer, multiple sclerosis, encephalitis, and diabetes [3]. Understanding biological barriers is also critical for drug development, as the absorption and distribution of pharmaceutical agents is affected by transport across barriers, determining drug efficacy and toxicity [1,4].

A key parameter for assessing *in vitro* models of biological barriers is barrier integrity. The classical method for evaluating endothelial barrier integrity is to use molecular tracer dyes (e.g., FITC-dextran) and monitor their diffusion across the monolayer over time. Electrical sensing provides a non-invasive alternative to molecular tracer dye assays that can interfere with molecular transport and barrier integrity [5]. Transepithelial/endothelial electrical resistance (TEER) is the gold standard metric used in non-invasive, label-free assessment of endothelial barrier integrity and permeability, and is an indicator of the conductance of ions across paracellular junctions [6]. This method provides the ohmic resistance of the monolayer. Major challenges for measuring TEER arise with the typical chopstick-like electrode handheld device format, with electrodes positioned on either side of a membrane insert. Using this format results in potential disruption of the cell layer through required culture device removal from incubator, as well as movement of electrodes from well to well [7]. Moreover, positioning of electrodes greatly impacts measured TEER values, as the electric field produced by the chopstick electrodes across the cell layer is not uniform [8]. Current distribution can also vary based on membrane surface area, electrode design, and device geometry, all of which can make it challenging to compare TEER values across platforms [9]. Increasing the size of the electrodes and integrating them into the measurement platform, as in the EndOhm system, reduces variability due to electric field non-uniformity and electrode positioning [10]. The TEER24 system (Applied Biophysics) further improves on this concept with the added functionality of continuous measurements for multiple wells over time, enabling non-invasive, real-time monitoring of biological barriers. However, due to their placement on the apical and basal side of the membrane, TEER electrodes do not directly capture the impedance of the monolayer; rather they measure the impedance of all material situated between the electrodes. In the context of co-culture setups, the presence of additional cell types and biomaterials (e.g., hydrogels) can confound TEER measurements.

An alternative method for electrically evaluating EC barriers is using impedance spectroscopy, which uses a range of measurement frequencies to differentiate between the contributions of the electrodes, bulk material and cell layer [11]. CellZScope incorporates this feature into a TEER-like setup of electrodes, with a measurement frequency range of 0-10^5^ Hz. Real-Time Cell Analysis (RTCA) from xCELLigence, on the other hand, allows culture of cells directly on electrodes, measuring high frequency responses (10 kHz, 25 kHz, 50 kHz). Electrical cell-substrate impedance sensing (ECIS), a technique developed by Keese and Giaever [12,13], combines culture of cells on electrodes with a range of frequencies, enabling direct measurement of the monolayer, with high sensitivity detection of nanometer-scale changes in cell junctions [12].

The format of ECIS has remained overall the same since its inception – cells are grown on gold electrodes integrated onto a solid substrate. Conventional ECIS shows a characteristic pattern for impedance based on measurement frequency. Low frequency impedance values are dominated by electrode capacitance, and high frequency impedance values by resistance of the culture medium [5]. Overall, mid-range frequencies (1-10 kHz) provide the greatest detection sensitivity [14,15], and cell spreading and proliferation is best monitored at higher frequencies (e.g., 40 kHz) [13], although the exact frequency values are cell type-dependent [14]. Although conventional ECIS offers many advantages, it does not allow for comparison against the gold-standard methods for electrical sensing of barrier integrity. Indeed, Bischoff et al. caution against directly comparing functional data from cells grown on permeable inserts vs. structural data of cells grown ECIS electrodes, as conventional ECIS electrodes are incorporated into a solid substrate, which does not allow for co-culture of cells relevant to the biological barrier [16].

To address this gap, we and others have integrated ECIS sensing capabilities into porous membranes configured as traditional Transwell-like cell culture inserts [17,18]. While these initial studies demonstrated the potential of porous membrane ECIS (PM-ECIS) to monitor the formation, disruption, and permeability of cell barriers, they did not systematically assess the effect of working electrode size, which is known to impact the dynamic range of ECIS measurements in traditional solid substrate systems [15,19–21]. Further, while PM-ECIS platforms enable chopstick TEER measurement, direct comparisons between PM-ECIS and TEER measurements and their electrode size dependency have not been made.

Here we assessed the effect of electrode size on PM-ECIS measurements of endothelial cell barrier growth and disruption and the correlation between PM-ECIS vs. TEER resistance. We found that PM-ECIS sensitivity to changes in barrier resistance and impedance depend on electrode size, with smaller electrodes being most sensitive to cell growth and barrier formation and disruption but only the largest electrodes showing positive correlation between PM-ECIS and TEER resistance readings. These data inform PM-ECIS device design and applications, and support its potential for direct, non-invasive, and real-time assessment of cells cultured on porous membranes.

## 2. Materials & Methods

### 2.1 Cell Culture

GFP-expressing HUVECs (AngioProteomie, cAP-0001GFP) were thawed and suspended in Endothelial Growth Medium (AngioProteomie, cAP-02). T75 flasks were pre-coated with 4 mL Quick Coating Solution (AngioProteomie, cAP-01) at room temperature for 5 min, then aspirated. HUVEC cell suspension (1×10^6^ cells) was transferred into the flask and placed overnight into a cell culture incubator (37°C, 5% CO_2_) to allow cells to adhere. Medium was replaced with 15 mL fresh EGM the next day, and changed every other day thereafter, and cells grown to confluence. HUVECs were cultured for 6-7 days prior to detachment with 0.25% trypsin-EDTA (Life Technologies, 25200-056) for use in experiments. Inserts were coated with bovine plasma fibronectin (Sigma-Aldrich, F1141) at 50 µg/mL, 2 mL in each insert, and cells (P5) seeded at a density of 80 000 cells/cm^2^ to achieve confluence by Day 1.

### 2.2 Device fabrication

Each device comprised gold electrodes patterned onto a porous transparent PETE membrane filter (3.0 µm, 6×10^5^ pores/cm^2^; Sterlitech, 1300032) on which endothelial cells were cultured. Device fabrication was performed based on published protocols [18]. Briefly, this process involves thermal bonding of gold leaf (Gold Leaf Supplies, 2300/RUXX) to the membrane, followed by UV lithography to pattern the electrodes, and laser cutting of the electrode-patterned membrane to size and shape (**Fig. 1A**).

**Fig. 1:**
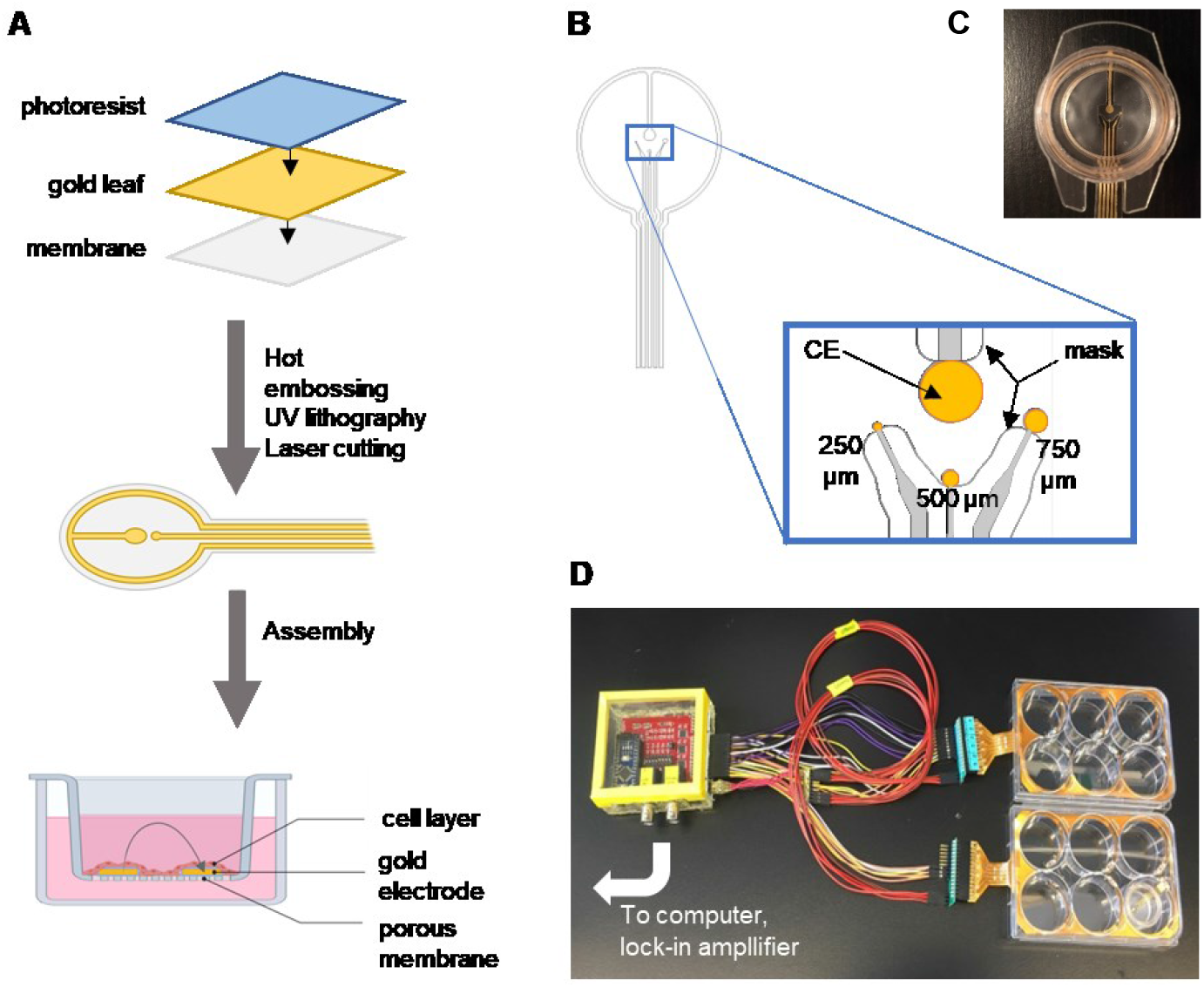
Fabrication and setup of 6-well ECIS membrane electrode devices. **A:** Fabrication process involves thermal bonding of gold leaf to membrane and UV lithography patterning of electrodes. The membrane electrodes are then assembled into a cell-culture insert format. Arrow in bottom schematic shows flow of current between ECIS electrodes through the endothelial cell layer and culture medium. **B:** Electrode design for testing three working electrode sizes of 250, 500, and 750 µm diameter. In the inset, gold denotes exposed electrode areas as defined by a tape mask; grey denotes electrode traces covered by the tape mask. Counter electrode = 2 mm diameter. Edge-to-edge working electrode-counter electrode distance = 1.5 mm. **C:** Top-down view of membrane electrode device. **D:** PM-ECIS devices were connected to a 24-channel multiplexer, which in turn connected to a lock-in amplifier and computer.

An electrode configuration was designed to test three different sizes of circular working electrode: 250 µm, 500 µm, and 750 µm diameters, with a circular counter electrode measuring 2 mm in diameter (**Fig. 1B, 1C**). Electrode diameters less than 250 µm were not tested, as only a few cells are sampled at that size and measurements are obscured by parasitic impedance from cables and measuring instruments at smaller diameters [19]. The spacing of each working electrode with respect to the counter electrode was 1.5 mm, measured by taking the shortest edge-to-edge distance between each working electrode and the counter electrode. The exposed area of the patterned electrodes was defined by a tape mask. The total insert area was 3.83 cm^2^.

### 2.3 ECIS & TEER measurements

The PM-ECIS devices were connected to a 24-channel multiplexer, which in turn connected to a phase-sensitive lock-in amplifier (LIA) and computer (**Fig. 1D**), The overall circuit for the ECIS system follows the set up described by Giaever and Keese [12]. The LIA acts to both generate an alternating current signal, and receive signal measured across the cell layer. An alternating current signal of 1 V rms was generated at a specified frequency, connected to a 1 MΩ resistor to provide a 1 µA current. Cell-free devices were filled with culture medium and measurements taken over at least 12 hours to establish baseline impedance. The voltage drop across the device containing cells grown on electrodes was detected and measured by the LIA, which allows for small AC signals to be resolved in high noise systems. A cell-free baseline was established beforehand using cell-free devices filled with culture medium and impedance measurements taken over the course of at least 12 hours prior to cell seeding. LIA outputs including voltage drop and phase angle were used to calculate and plot impedance and resistance values using a custom MATLAB program [18].

TEER was measured using a Millicell ERS volt-ohm meter (World Precision Instruments, New Haven, CT) according to the manufacturer’s instructions.

### 2.4 Immunostaining

Cells were fixed in methanol for 10 min at −20°C, washed three times with PBS, and blocked with 3% BSA (Sigma-Aldrich, 10735086001) for 20 min at 37°C. Cells were stained with primary antibodies (1:100) diluted in 3% bovine serum albumin in PBS for 1 hour at 37°C. Following incubation, cells were washed three times with PBS and blocked using 10% goat serum (Sigma-Aldrich, G9023) in PBS for 1 hour at room temperature. Secondary antibodies (1:200) diluted in 10% goat serum were applied for 1 hour at room temperature. After washing three times with PBS, nuclei were stained with 1:1000 Hoechst 33342 (Sigma-Aldrich, B2261) for 5 min at room temperature. Primary antibodies were rabbit polyclonal VE-cadherin (Abcam, ab33168) and mouse monoclonal ZO-1 (Thermo Fisher Scientific, 33-9100); secondary antibodies were goat anti-mouse IgG (H+L) Alexa Fluor 647 (Life Technologies, A-21236) and goat-anti rabbit IgG (H+L) Alexa Fluor 568 (Thermo Fisher Scientific, A-11011). Image acquisition was performed using an Olympus FV3000 confocal laser scanning microscope or Olympus BX51 inverted fluorescence microscope as indicated.

### 2.5 EGTA treatment assay

Cells were incubated in 5 mM ethylene glycol-bis(β-aminoethyl ether)-N,N,N′,N′-tetraacetic acid EGTA (Sigma-Aldrich, E3889) diluted in Endothelial Growth Medium at 37°C and 5% CO_2_ for 2 hours, with ECIS measurements taken every 15 min. After 2 hours, EGTA was removed and replaced with fresh growth medium.

### 2.6 NaCl concentration curve

Cell-free PM-ECIS 6-well inserts were filled with varying concentrations of NaCl (0.005, 0.01, 0.1, 1 M) and measurements taken for each concentration by chopstick TEER and ECIS. Values reported for TEER are resistance as measured by the Millicell ERS volt-ohm meter without adjustment. ECIS values are resistance values measured without adjustment.

### 2.7 Comparison TEER vs. ECIS calculations

TEER values were calculated using TEER = (R_cell_ – R_cell-free_) x surface area of insert. ECIS values were calculated using ΔR = R_cell_ – R_cell-free_. TEER and ECIS values were normalized to the 6 h time point for each of the experimental runs, with up to three devices tested per run.

## 3. Results and discussion

### 3.1 Characterization of cells grown on ECIS membrane device

To assess the ability of endothelial cells to form confluent monolayer on different sizes of PM-ECIS working electrodes, GFP-expressing HUVECs were seeded at 80 000 cells/cm^2^ and cultured for five days in devices with one counter electrode and three different working electrode sizes of 250, 500, and 750 μm diameter. Each working electrode measured its corresponding cell population independently from the other working electrodes in the same device. Immunostaining confirmed that the PM-ECIS devices supported the growth of confluent HUVEC monolayers, with VE-cadherin continuously distributed around the cell peripheries (**Fig. 2A**) and punctate ZO-1 membrane expression across the cell layer (**Fig. 2B**), consistent with previous observations of HUVEC monolayers on tissue culture-treated polystyrene [22] and porous membrane [23] substrates. Notably, the HUVEC monolayer spanned the electrode-membrane boundary with no visible effect on morphology or expression of junctional proteins compared to surrounding cells (**Fig. 2C**).

**Fig. 2:**
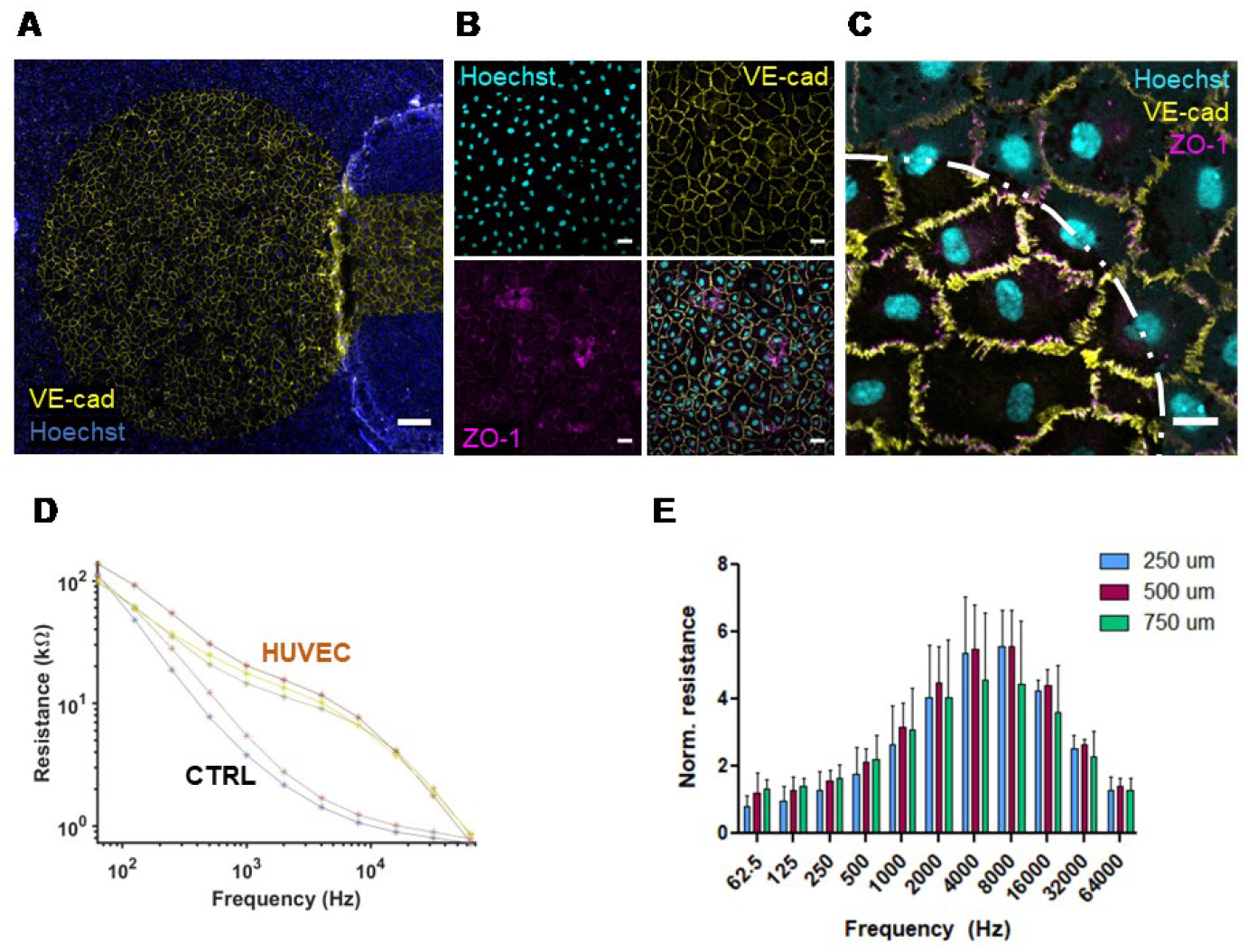
ECIS membrane device supports growth of confluent endothelial monolayer. **A:** Fluorescence microscopy images of human umbilical vein endothelial cells (HUVECs) covering counter electrode, stained for VE-cadherin (yellow) and nuclei visualized with Hoechst 33342 (blue). Scale bar = 200 µm. **B:** Confocal microscopy imaging shows expression of VE-cadherin (yellow) and ZO-1 (magenta) in HUVECs grown on electrode surface. Nuclei visualized with Hoechst 33342. Scale bars = 50 µm. **C:** Confocal microscopy imaging shows HUVEC monolayer spanning the electrode-membrane boundary. Scale bar = 20 µm. **D:** Representative frequency vs. resistance plot for Day 4 post-confluence HUVECs grown on 250 µm electrodes; CTRL = cell-free control. Each line corresponds to a replicate in a separate cell culture insert. **E:** Normalized resistance values across the scanned frequencies. Normalized resistance (R_cell_/R_cell-free_) for three electrode sizes (d= 250, 500, 750 µm) across frequencies. N = 5-7 cell-covered, 6-7 cell-free for each electrode size.

To validate PM-ECIS vs. conventional ECIS measurements in the literature, frequency scans were performed on HUVECs four days post-confluence, with measurements taken at f = 62.5, 125, 250, 500, 1 000, 2 000, 4 000, 8 000, 16 000, 32 000, and 64 000 Hz. PM-ECIS yielded the expected frequency-resistance relationship for cell-covered vs. cell-free electrodes, with the cell layer contributing to a characteristic increase in signal over the mid-range frequencies compared to cell-free electrodes [19,24] (**Fig. 2D**). Normalized resistance values of HUVECs on PM-ECIS devices peaked at ∼4 kHz for all electrode sizes, consistent with the 4 kHz measurement frequency conventionally used for HUVEC monolayers in solid substrate ECIS setups [19] (**Fig. 2E**).

### 3.2 Effect of electrode size on PM-ECIS measurements of monolayer growth and barrier integrity

The contribution to overall impedance by the electrode-solution interface is proportional to 1/r^2^, where r is radius of the working electrode; thus, impedance is inversely proportional to electrode size, with smaller working electrodes expected to be more sensitive than larger ones [19]. To test this in PM-ECIS devices, growth of HUVECs was monitored on three electrode sizes (d = 250, 500, 750 μm) over 5 days. To assess electrode size effects on PM-ECIS measurement of barrier formation, change in normalized resistance at 4 kHz was measured throughout the culture period and compared between electrode sizes. Resistance increased over the first ∼2 days of culture on all electrode sizes, with the change in resistance being inversely proportional to the electrode size (**Fig. 3A**). From Day 3 to 5 of culture, resistance values plateaued for all electrode sizes, indicating formed barriers (**Fig. 3A**). Endpoint analysis on Day 5 confirmed a significant effect of electrode size on normalized resistance (p<0.001), with the greatest change on the smallest (250 µm) electrodes (**Fig. 3B**). To evaluate whether the same trends applied to cell spreading and proliferation across the PM-ECIS electrodes, impedance was measured at 40 kHz. Indeed, there was a significant effect of electrode size on normalized impedance (p<0.001), with the greatest change on the smallest electrodes (**Fig. 3C**).

**Fig. 3:**
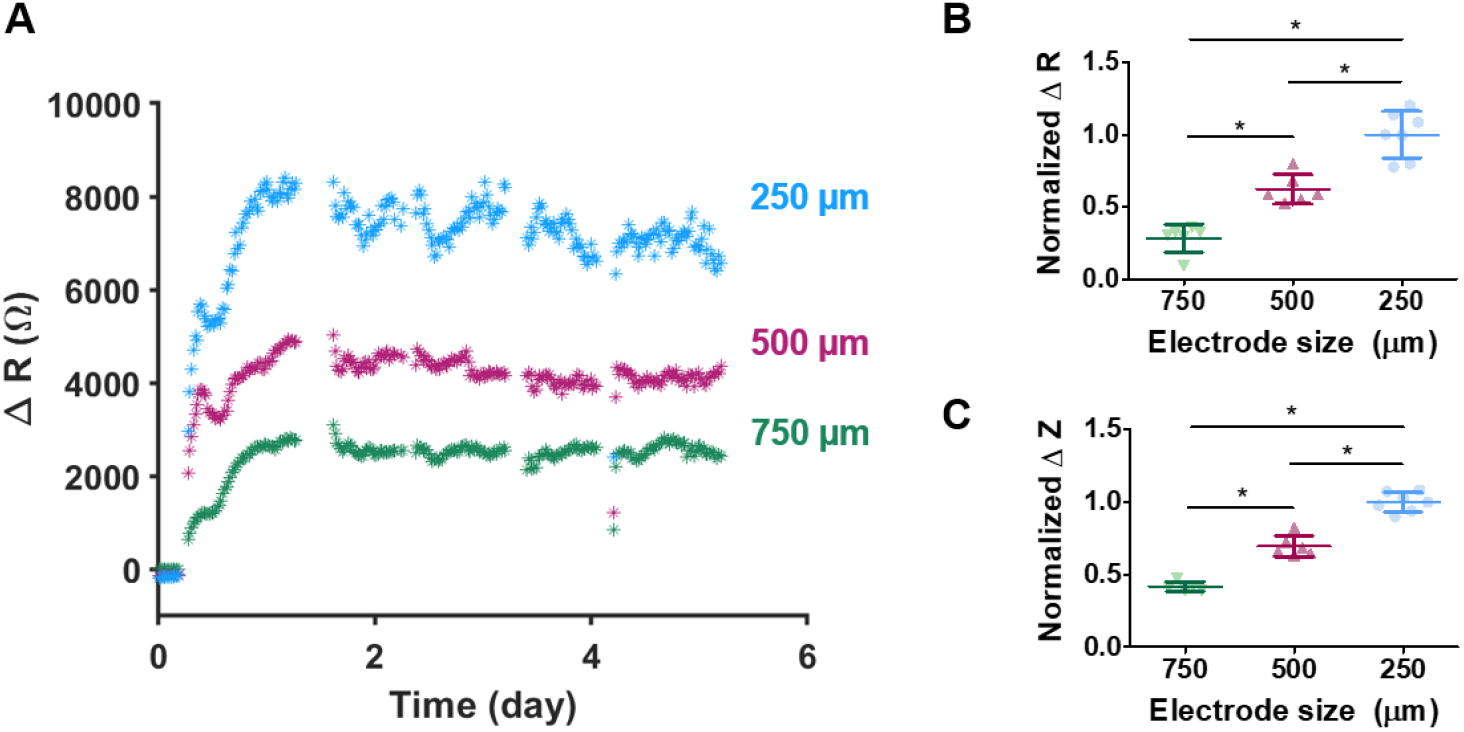
Electrode size-dependent response in impedance and resistance for HUVEC monolayer grown in PM-ECIS devices. **A:** Representative ΔR (4 kHz) vs. time plot for Day 4 post-confluence HUVECs. ΔR = R_cell_ – R_cell-free_ for corresponding electrode sizes. **B, C:** Normalized R and Z values for Day 4-post confluence HUVECs grown on three electrode sizes (d= 250, 500, 750 µm) **B:** ΔR normalized to 250 µm electrodes for measurements taken at 4 kHz, and **C:** ΔZ normalized to 250 µm electrodes for measurements taken at 40 kHz. ΔZ = Z_cell_ – Z_cell-free_ for corresponding electrode sizes. N = 5-7 cell-covered and 6-7 cell-free electrodes for each electrode size. \**p*<0.001 using one-way ANOVA with Bonferroni test.

These trends on PM-ECIS devices are consistent with those reported by others on traditional ECIS. For example, a study that measured impedance of confluent bovine aortic endothelial cells on solid substrate ECIS working electrodes with size ranging from d = 200 to d = 800 µm, experimentally demonstrated a relationship between smaller electrodes and greater impedance values [20]. The same study showed that an increase in cell density corresponded to a greater increase in impedance on working electrodes with d = 200 µm compared to d = 300 µm. Experimentally, however, electrodes smaller than d = 250 µm may not be beneficial. One study used four different adherent cell types to compare the peak value of normalized resistance on frequency scans for various electrode sizes (d = 25, 50, 100 and 250 μm) [25]. The authors found that this peak value – which represents the maximal relative change in resistance contributed by the cells – decreased with electrode size, with the greatest peak value for confluent monolayers being measured with the 250 µm electrodes. Moreover, the authors saw the highest apparent peak in normalized resistance using d = 250 µm when measuring cell response to TGF-β1 treatment, and no significant cell response detected by electrodes smaller than d = 100 µm, suggesting that statistical significance of cell response is related to the size of the cell population being measured [25].

### 3.3 Measurement of response to EGTA barrier disruption

To assess the effect of electrode size in PM-ECIS response to barrier disruption, a confluent monolayer of HUVECs was subjected to 2-hour treatment with EGTA. EGTA disrupts cell barriers by chelating Ca^2+^ ions to destabilize tight junctions [26,27]. Therefore, a decrease in barrier integrity, and by extension, ECIS resistance, would be expected after EGTA treatment. Barrier disruption with EGTA treatment resulted in a drop in resistance within 15 minutes, measured at 4 kHz for all electrode sizes (**Fig. 4A**; p<0.01 for 250 and 750 µm electrodes and p=0.0575 for 500 µm electrodes). While all electrodes sizes were sensitive to barrier disruption, the reduction in barrier resistance was significantly greater with the 250 µm (p<0.001) and 500 µm (p<0.05) electrodes than the 750 µm electrodes (**Fig. 4B**).

**Fig. 4:**
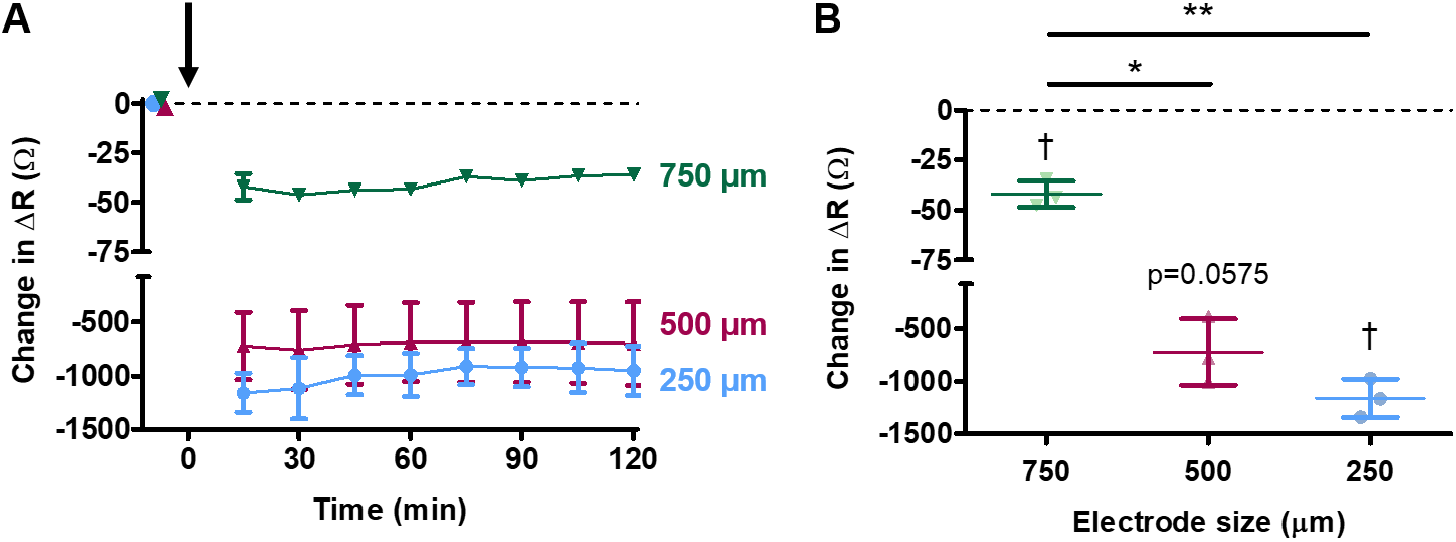
Electrode size-dependent response in impedance and resistance to barrier disruption with EGTA. A: Change over time in ΔR from baseline at 4 kHz after treatment with EGTA in confluent HUVEC monolayers. ΔR = Rcell – Rcell-free; Change in ΔR = ΔR(t=0) – ΔR(t) for corresponding electrode sizes. Black arrow indicates time that EGTA was added. B: Change in ΔR after 2 h EGTA treatment. N = 3-4 cell-covered and 3-4 cell-free electrodes for each electrode size. †p<0.01 by one-sample t-test relative to change in ΔR=0; * *p* < 0.05, ** *p* < 0.01 by one-way ANOVA with Bonferroni test.

### 3.4 Comparison of PM-ECIS to TEER

To further validate PM-ECIS, comparison to chopstick TEER was initially performed using NaCl concentration curve measurements, a method previously used by Cacopardo et al. for impedimetric system assessment [1]. Cell-free PM-ECIS 6-well inserts were filled with varying concentrations of NaCl (0.005, 0.01, 0.1, 1 M) and resistance measured by chopstick TEER and ECIS. As expected, the resistance measured by both TEER and PM-ECIS was NaCl concentration-dependent, with statistically significant correlation between the two methods for all three electrode sizes (**Fig. 5**; r>0.9; p<0.0001).

**Fig. 5:**
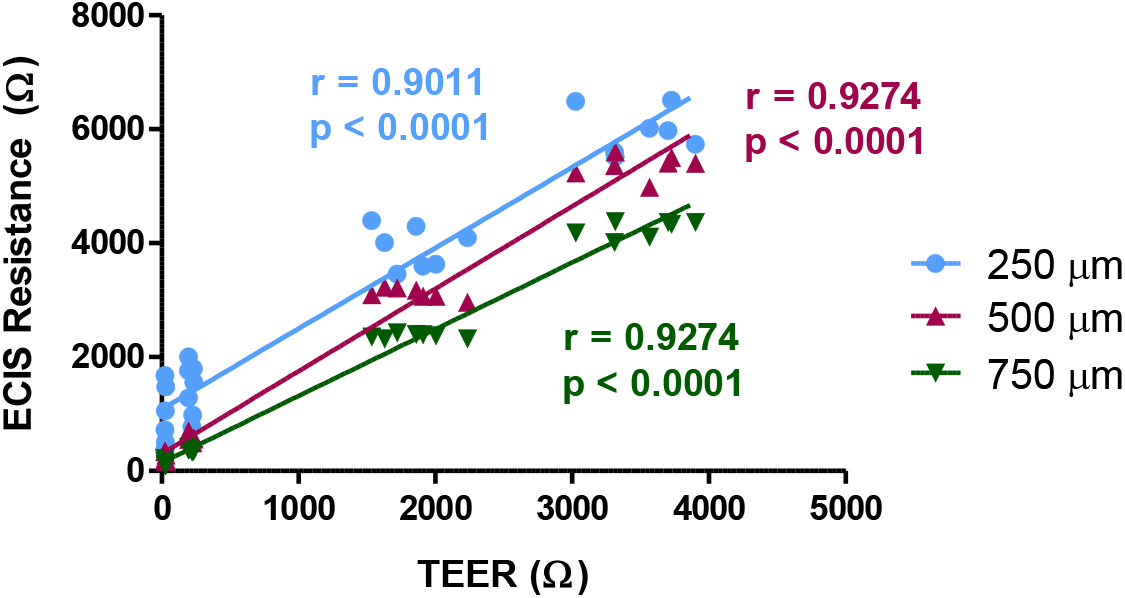
Comparison of ECIS membrane method vs. TEER. **A:** Correlation between 4 kHz ECIS resistance and TEER resistance across varying concentrations of NaCl solution (0.005, 0.01, 0.1, 1M) for 250, 500, and 750 μm working electrodes. Each data point represents mean for N= 7-8 electrodes. Pearson correlation analysis with trendlines from linear regression for illustrative purposes.

Having shown strong correlation between ECIS and TEER for NaCl solutions, correlation between the two methods for endothelial cell monolayers was next assessed. HUVECs were seeded in devices with all three electrode sizes, and normalized TEER and ECIS measurements were taken over 48 h post-seeding. ECIS measurements from the 250 and 500 µm diameter electrodes were not correlated to TEER values obtained for same cell monolayer (**Fig. 6A, 6B**; p=0.46 and p=0.39, respectively). This likely reflects differences in the spatial scales to which the two methods are sensitive: TEER measures resistance across the entire cell monolayer and media compartments, whereas small PM-ECIS electrodes sample a smaller number of cells with high sensitivity to local changes in the cell monolayer (**Figs. 3, 4**). Consistent with this, normalized ECIS resistance for HUVECs grown on the largest 750 µm electrodes was positively and significantly correlated to normalized chopstick TEER values (**Fig. 6C;** p = 0.058).

**Fig. 6:**
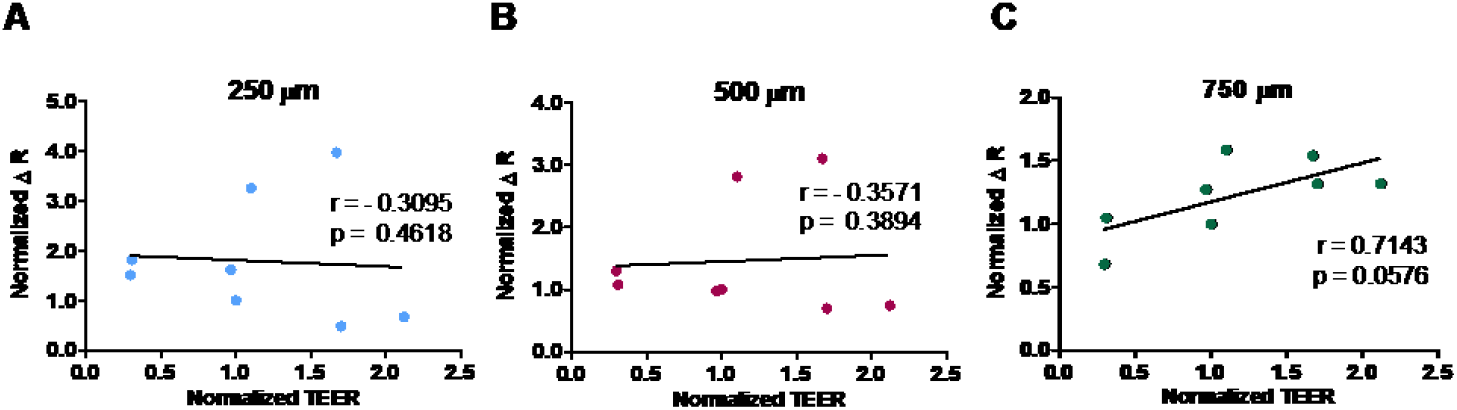
Comparison of ECIS membrane method vs. TEER in HUVEC monolayers. **A, B, C:** Correlation between normalized TEER and ΔR for HUVEC across range of timepoints (6, 24, 48 h). ΔR = R_cell_ – R_cell-free_; N = 2-3 cell-covered and 6-8 cell-free electrodes per timepoint, for each electrode size. ΔR and TEER normalized to 6 h time point. **A:** 250 μm working electrodes. **B:** 500 μm working electrodes. **C:** 750 μm working electrodes. Pearson correlation analysis with trendlines from linear regression for illustrative purposes.

The trade-off between signal sensitivity and cell population sampled with small single working electrodes could be addressed with multiple small electrodes measuring in parallel to a common counter electrode. Such a design would increase the sampling area albeit with a small reduction in sensitivity compared to a single working electrode for confluent monolayers normalized to cell-free electrodes [19]. Single electrodes do, however, provide a higher magnitude signal in response to, and better resolution of slope differences over, the various stages of cell adhesion and movement when measuring rate of change in impedance over time [28]. This suggests a particular relevance of the single electrode configuration tested in this study to assays where time resolution of cell response to a treatment condition is being evaluated.

## 4. Conclusions

In this study, PM-ECIS was validated across three different working electrode sizes (d = 250, 500 and 750 µm). An electrode size-dependent response was demonstrated for monolayer growth, barrier formation, and barrier disruption with EGTA. Importantly, this study showed a direct correlation between PM-ECIS and TEER measurements, indicating its potential to replace conventional TEER due to its capacity for direct, non-invasive, and real-time assessment of cells cultured on porous membrane, with tunable sensitivity based on electrode size. Notably, the electrode dimensions and fabrication method for PM-ECIS make it compatible not only with conventional co-culture setups, but also with microfluidic organ-on-a-chip configurations.

## CRediT authorship contribution statement

**Alisa Ugodnikov:** Conceptualization, Methodology, Formal analysis, Investigation, Data curation, Writing – Original draft, Project administration. **Oleg Chebotarev**: Methodology, Software, Funding acquisition. **Henrik Persson**: Investigation. **Craig A. Simmons**: Conceptualization, Methodology, Resources, Writing – review, Supervision, Project administration, Funding acquisition.

## Acknowledgments

This project was funded by the Natural Sciences and Engineering Research Council of Canada (CRDPJ 531083-18 and CREATE Training Program in Organ-on-a-Chip Engineering and Entrepreneurship) and the Ontario Centre of Excellence (VIP II 29253).

